## Supplementary Information for "Programmable Viscoelasticity in Protein-RNA Condensates with Disordered Sticker-Spacer Polypeptides"

*All correspondence could be addressed to:

### Methods

**Peptide and RNA samples.** See Table-S1 for a list of peptides used in this study. All peptides ($[RGRGG]_5$, $[KGKGG]_5$, $[RGYGG]_5$, $[RGPGG]_5$, $[RGFGG]_5$, $[RGSGG]_5$, $[RARAA]_5$, $[RQRQQ]_5$, $[RPRPP]_5$, $[RLRLL]_5$, RG1Y, RG3Y, FUS-RGG, TAF15-RGG, and EWS-RGG were synthesized by GenScript USA Inc. (NJ, USA, >90% purity). All peptides contained a cysteine at the C-terminal for site-specific fluorescence labeling. Peptides were reconstituted in RNase-free water (Santa Cruz Biotechnology) containing 50 mM dithiothreitol (DTT) to prevent cysteine oxidation and the stock solutions were split into multiple aliquots and stored at -20 $^{o}$C. Homopolymeric RNA $[rU]_{40}$ (MW 12,185 Da), ssDNA $[dT]_{40}$ (MW 12106 Da), $G_5T_{30}C_5$ and $(G4C2)_5$ ssNA were purchased from Integrated DNA Technologies (IDT) and reconstituted in RNase-free water. All peptide and RNA stock solutions were checked under the microscope to ensure complete solubilization and the absence of aggregates. No additional purification was performed on the peptides or the RNA.

**Peptide-RNA condensate preparation.** Each type of condensate was prepared by mixing peptide and RNA in a buffer containing 25 mM Tris-HCl (pH 7.5), 25 mM NaCl and 20 mM DTT. The peptide final concentration was 5.0 mg/ml and the rU40 RNA concentration was 2.5 mg/ml, corresponding to a mass ratio of 0.5. This ratio was chosen based on solution turbidity measurements of $[RGRGG]_5$-rU40 mixtures which showed maximum absorbance[1] (indicating a maximum degree of condensation; Fig. S14a). All peptides underwent LLPS with RNA at this ratio and resulted in the formation of condensates which were then subjected to microrheology experiments to measure their linear viscoelastic behavior. Due to the comparative nature of this study, all buffer conditions were kept identical. For microrheology experiments, yellow-green carboxylate-modified polystyrene beads (1 µm in diameter, FluoSpheres™, Invitrogen) were added to the buffer at a low concentration (0.0003 % solids). These beads were found to partition well within peptide-RNA condensates[1]. For experiments performed with $G_5T_{30}C_5$ and $(G4C2)_5$ ssNAs, the same buffer was used except for the addition of 2.5 mM $MgCl_2$. For FET-RGG condensates (TAF15, EWS, and FUS), the polypeptide concentration was 10 mg/ml, the ssDNA T40 concentration was 1 mg/ml, and the buffer contained 25 mM Tris-HCl (pH 7.5), 25 mM NaCl and 20 mM DTT.

**Passive microrheology with optical tweezers (pMOT).** The method of passive microrheology used in our study was introduced by Tassieri, Preece, and Evans[2-5] and is built on the work on optical tweezer-based microrheology by Mason and Weitz[6]. The complex modulus of a material (condensate) is calculated from the analysis of the position fluctuations of an embedded probe particle trapped with an optical tweezer. The motion of a trapped bead within the medium is mediated by the thermal fluctuations of the medium and constrained by the harmonic potential of the optical trap. This is one of the few microrheology methods that do not require an independent optical trap calibration procedure[5].

Condensate-forming samples (5 µl) of peptides and RNA were placed on a tween20-coated microscope coverslip (0.17 mm thickness, 18x18 mm dimensions). Next, a glass slide with two double-sided tape strips was gently placed on the sample, sandwiching it between the coverslip and the glass slide. ~100 µl of mineral oil was injected into the chamber between the coverslip and the glass slide such that oil surrounds the sample from all directions. This was done to ensure that the sample does not experience evaporation or evaporation-induced flow. The condensate sample was left to equilibrate for a minimum of 1 hour. The pMOT experiments were not initiated until all droplets were settled on the coverslip surface and no droplet fusion events were observed. To measure the frequency-dependent viscoelastic moduli, the custom-made flow chamber was loaded onto a correlative confocal microscope-optical tweezer setup (LUMICKS C-Trap) with a 60x water immersion objective and a bright-field camera. Next, the optical trap was used to trap a bead that is embedded within a peptide-RNA condensate. The trapping power was set to ~100 μW initially to trap the bead. Using the optical trap, the bead was then positioned at the center of the condensate and ~2-5 μm above the coverslip surface. The trapping power was lowered to the minimum value that ensures constraining the bead within the optical trap. The trap-constrained motion of the bead was tracked using the bright field camera with a 500 Hz acquisition rate. The trajectory of the bead (X-Y coordinates as a function of time) was extracted using the built-in tracking algorithm in the instrument (LUMICKS C-trap). For each peptide-RNA system, we collected trajectories from 3-5 condensates for three independently prepared samples (a total of 9-15 trajectories). Every single trajectory was obtained by tracking the bead for a minimum of 10 minutes and a maximum of 45 minutes depending on the peptide-RNA system and the quality of the extracted autocorrelation curves (see the data analysis section). All trajectories were analyzed using custom-build python scripts to obtain frequency-dependent viscoelastic moduli and viscosity. Trajectories that showed asymmetric motion in the X-Y direction were discarded. The analysis of the pMOT experiments as well as additional control measurements are discussed in detail in Supplementary Note-1.

**Video particle tracking microrheology (VPT).** 200 nm yellow-green carboxylate modified polystyrene beads (FluoSpheres, Invitrogen) were used for VPT measurements. The samples were prepared at identical conditions as described in the sample preparation section. Fluorescence video imaging was done using a Zeiss Primovert inverted microscope equipped with a 100x oil immersion objective and a Zeiss axiocam 503 monochrome camera. Movies of the beads diffusing within the condensate were collected for approximately 2-5 minutes. Particle tracking was performed using Trackmate[7] plugin in Fiji-ImageJ. Mean squared displacement (MSD) was calculated from the trajectories for several lag times $\tau$ using home-built python scripts. The ensemble average MSD was calculated (from ~20-100 individual particles) and fitted with $MSD(\tau) = 4D\tau^{\alpha} + b$. In the previous equation, D is the diffusion coefficient, $\alpha$ is the diffusivity exponent and b is a constant accounting for the noise. The condensate viscosity $\eta$ was calculated from the Stokes-Einstein equation[1]

$$\eta = \frac{k_B T}{6\pi D a} \tag{1}$$

Where $k_B$ is Boltzmann constant, $T$ is the temperature, and $a$ is the particle radius. For each condensate type, we collected trajectories from 3-5 condensates in 3 independently prepared samples. Each condensate contained somewhere between 20-100 microspheres. Trajectories of beads close to the condensate surface were excluded from the MSD calculation. For $[RGYGG]_5$-rU40 and $[RGRGG]_5$-rU40 condensates, the MSD was fitted with $MSD = d^2(1 + \frac{\tau}{\tau_c})$, which is the predicted behavior of a Maxwell fluid[8].

**Fluorescence recovery after photobleaching (FRAP).** Peptide-RNA condensates were prepared using 5.0 mg/ml peptide and 2.5 mg/ml rU40 RNA mixed in 25 mM Tris-HCl (pH 7.5), 25 mM NaCl, 20 mM DTT containing ~500 nM of FAM-labeled RNA oligo rU10 (purchased from IDT). Samples were injected in a Tween20-coated imaging chamber and loaded to the confocal microscope stage (LUMICKS C-trap). Bleaching was achieved using a 488 nm laser at 100% laser power for ~0.5 seconds. The fluorescence intensity of a given region of interest (ROI) was then recorded until the recovery was complete. The bleaching ROI size and shape were kept identical across all samples. The intensity time traces were plotted for comparison and extraction of $t_{1/2}$ values. For each peptide-RNA sample, 5 recovery traces were collected and averaged. Intensity error bars were estimated using the standard deviation at each recorded time point.

**Temperature-Salt state diagram measurements:** Peptide-RNA condensates were prepared using 5.0 mg/ml peptide, 2.5 mg/ml rU40 RNA mixed in a buffer (25 mM Tris-HCl, pH 7.5 and 20 mM DTT) with the desired salt concentration as noted in Figure 4 in the main text. Peptide-RNA samples were then sandwiched between a PEG5000-coated glass slide and a coverslip. The sample was surrounded by mineral oil to prevent sample evaporation due to heating. The sample was then placed into a custom-built temperature stage (Instec Inc., temperature range: 5-90 $^{o}$C) and loaded on a Zeiss Primovert inverted microscope equipped with a 40x objective and a Zeiss axiocam 503 monochrome camera. To determine the phase separation temperature ($T_{ph}$), the sample was heated first to maximum temperature (90 $^{o}$C) until all condensates were dissolved. Next, the sample was cooled down in steps of 5 $^{o}$C and an equilibration time of 5 minutes until LLPS was observed. Once the LLPS transition was noted, samples were heated again until condensates were dissolved and the LLPS temperature was subsequently approached in steps of 2 $^{o}$C and equilibration time of 5 minutes in each temperature to determine the $T_{ph}$.

**Mesh size determination experiments:** Peptide-RNA samples were prepared at 5.0 mg/ml peptide and 2.5 mg/ml rU40 RNA in a buffer containing 25 mM Tris-HCl (pH 7.5), 25 mM NaCl and 20 mM DTT. After mixing the peptide and the RNA and the formation of condensates, a small concentration (~500 nM) of tetramethylrhodamine-labeled dextran of the desired size was added to the sample. The sample was mixed and subsequently imaged under a confocal microscope (LUMICKS C-trap). If the hydrodynamic radius of the dextran molecule is smaller than the mesh size of the condensate, it is expected that dextran molecules will be positively recruited into the condensates. However, if the hydrodynamic radius of the dextran molecule is larger than the mesh size of the condensates, dextran recruitment will not occur. The partition behavior of dextrans of variable molecular weights (and variable hydrodynamic radii) was recorded and the mesh size range was determined as previously reported[9,10].

**Glass slide and coverslip preparation:** For the temperature-salt state diagram, mPEG5K-Silane was used to coat glass slides and coverslips to prevent condensates from sticking to the glass surfaces. The PEG coating procedure was performed according to the description of Alberti and coworkers[11]. Briefly, Glass slides were incubated in 2% Hellmanex solution for two hours and then rinsed with milliQ water and dried under compressed airflow. Next, the glass slides were immersed in a solution containing 20 mg/ml of methyl-PEG5K-Silane and 100 mM HCl dissolved in toluene and incubated for 18 hours in a closed glass staining jar. Glass slides were then rinsed with toluene, 90% ethanol, and MilliQ and dried under compressed airflow.

For all other experiments in this work, Tween20 coating was used to prevent droplet spreading on the glass surface. Glass slides and coverslips were first cleaned with 70% ethanol and dried under compressed airflow. Next, glass slides and coverslips were immersed in a 20% vol/vol solution of Tween20 for 30 minutes. Subsequently, glass slides and coverslips were rinsed 6-7 times with MilliQ water and dried using compressed air. Finally, glass slides and coverslips were dried in an oven set at 40 $^{o}$C for 4-8 hours and stored at room temperature for later use.

**Bead Halo assay:** Samples were prepared by mixing $[RGRGG]_5$ with rU40 RNA at 5.0 mg/ml peptide and 2.5 mg/ml RNA concentrations in a buffer containing 25 mM Tris-HCl (pH 7.5), 25 mM NaCl, and 20 mM DTT. Carboxylate-coated beads were added to the buffer along with the bait molecule (either $[RGRGG]_5$-A594 or rU10-FAM) before mixing the peptide and RNA. The bait molecule concentration was kept at ~200 nM. Upon mixing the peptide and the RNA, condensates were formed. The sample was thoroughly mixed and then sandwiched between a coverslip and a microscope glass slide spaced by two layers of double-sided tape. The sample was kept to equilibrate for 10 minutes and then loaded onto a confocal microscope (LUMICKS, C-trap). Several bead-harboring condensates were imaged using both brightfield and fluorescence illumination. For each bait molecule ($[RGRGG]_5$ or rU10), two independent samples were prepared and several condensates were imaged. Intensity profiles were plotted to inspect the fluorescence intensity at the bead surface and compare it with the intensity inside the droplet.

**All-atom simulations:** The explicit all-atom (AA) simulations with model tri-nucleotide and tri-peptide systems were done with OpenMM 7.5 using Amber force field a99SB-disp*-ildn force field with TIP3P model for water. The system consisted of tri-peptides and tri-nucleotides packed in a cubic box with concentration ~100 mg/ml, solvated, and charge neutralized with NaCl ions at 10mM concentration at peptide:RNA ratio of 1:2. The total number of atoms was ~100,000 atoms. 10 independent simulations were run for 1 microsecond. The combined trajectory from the 10 replicas was used for deriving the potentials of mean force. For each simulation trajectory, integration time was set to 2 fs. After minimization, the simulation was done in an NPT ensemble where the pressure was kept at 1 atm via Monte Carlo Barostat and the temperature was kept at 300K using a standard Langevin integrator with 1/ps friction coefficient.

**Phase coexistence simulations:** The simulations with coarse-grained (CG) models of protein and nucleic acid chain mixtures were done using HOOMD-Blue 2.9.6 molecular dynamics library. We used a 1 bead per residue model for amino acids and RNA based on formulation first presented in Alshareedah and coworkers[9]. The short-range potentials were calibrated according to the all-atom simulations to account for the differential stickiness of amino acids. Slabs were made by first randomly packing polypeptide and RNA chains in a cubic box. The box was then compressed to ~25 nm over the course of $10^5$ steps at a high temperature corresponding to T=600. The z-axis of the box was then extended to ~125 nm, and the packing was equilibrated for $10^6$ steps to allow the dense and dilute phases to form while quenching the system to the target temperature. Finally, the equilibrated dense and dilute phases were simulated for an additional $10^7$ steps to collect data. Simulations were run at various temperatures with 4000 polypeptide chains and an appropriate number of RNA chains ensuring a net chain charge neutrality. In all simulation steps, a Langevin integrator was used with a friction coefficient of 0.01.

**Bioinformatics analysis:** Amino acid sequences of RNA-binding proteins were downloaded from Uniprot (RefSeq) based on the annotation provided in RBPDB[12]. RG/RGG motifs within these proteins were extracted using a custom python script. RG/RGG motif was defined as per Thandapani *et al.*[13]. Briefly, peptide motifs with the sequences RG-$x_{(0\text{-}4)}$-RG and RGG-$x_{(0\text{-}4)}$-RGG were extracted, where *x* can be any amino acid. A binary presence/absence test was performed to determine how many of the extracted sequences contain each of the 20 amino acids within the spacer [*i.e.,* $x_{(0\text{-}4)}$]. Results are presented as a bubble chart in the main text Fig. 2c, where the diameter of an individual bubble corresponds to the fraction of spacers that contain respective amino acids. Cyan bubbles represent fractions > 10%.

**Software:** pMOT analysis was done using custom-made python scripts. Fiji-ImageJ[14] (version 1.52p) was used for image processing. OriginPro (2018b) was used for Graphing. Adobe Illustrator CC (2019, v23.0) was used for the figure assembly and production. ZEN (blue, v2.3) was used for image recording using a Zeiss Primovert microscope. Bluelake (v1.6.11) was used for image recording and particle tracking using Lumicks C-Trap microscope. HOOMD-Blue 2.9.6

was used for coarse-grained molecular dynamics simulations and OpenMM 7.5 was used for all-atom simulations. Ovito 3.5 was used for generating phase coexistence snapshots and ngvliew 3.0 was used for visualizing all-atom residue-base interaction snapshots shown in Figure 4.

**Statistics and reproducibility.** All experiments reported in this manuscript have been repeated at least three times with consistent results. The viscosity measurements are repeated for at least 12 data points from different condensates across three independently prepared samples. The relaxation times are obtained from at least 12 data points from different condensates across three independent sample preparations. The VPT experiments were replicated three times across different sample preparations. Bright-field and fluorescence microscopy images are representative of a large set of images exhibiting similar features. For temperature-salt state diagrams, random points were selected and reproduced three times successfully. FRAP experiments were repeated five times (within 1-2 sample preparations) for each peptide-RNA system.

**Table S1.** Amino acid sequences of the peptides used in this study.

| Peptide name | Sequence |
|---|---|
| $[RGRGG]_5$ and RG0Y | RGRGG RGRGG RGRGG RGRGG RGRGGC |
| $[KGKGG]_5$ | KGKGG KGKGG KGKGG KGKGG KGKGGC |
| $[RGFGG]_5$ | RGFGG RGFGG RGFGG RGFGG RGFGGC |
| $[RGYGG]_5$ and RG5Y | RGYGG RGYGG RGYGG RGYGG RGYGGC |
| $[RGSGG]_5$ | RGSGG RGSGG RGSGG RGSGG RGSGGC |
| $[RGPGG]_5$ | RGPGG RGPGG RGPGG RGPGG RGPGGC |
| $[RPRPP]_5$ | RPRPP RPRPP RPRPP RPRPP RPRPPC |
| $[RARAA]_5$ | RARAA RARAA RARAA RARAA RARAAC |
| $[RQRQQ]_5$ | RQRQQ RQRQQ RQRQQ RQRQQ RQRQQC |
| $[RLRLL]_5$ | RLRLL RLRLL RLRLL RLRLL RLRLLC |
| RG1Y | RGRGG RGRGG RGYGG RGRGG RGRGGC |
| RG3Y | RGRGG RGYGG RGYGG RGYGG RGRGGC |
| TAF15-RGG box | RGYGGDRGYGGDRGYGGDRGYGGDRGYGGDC |
| EWS1-RGG box | RGRGGPGGMRGGRGGLMDRGGPGGMFRGGRGGC |
| FUS-RGG box | RRGGRGGYDRGGYRGRGGDRGGFRGGRGGGDRGC |

**Table S2.** Viscosity and terminal relaxation time of the various peptide-RNA condensates tested in this study.

| Peptide-RNA | Viscosity (Pa.s) | Terminal relaxation time (ms) |
|---|---|---|
| $[RGRGG]_5$ and RG0Y | 4 ± 1 | 60 ± 10 |
| $[KGKGG]_5$ | 0.27 ± 0.05 | N/A |
| $[RGFGG]_5$ | 10 ± 3 | 110 ± 40 |
| $[RGYGG]_5$ and RG5Y | 40 ± 10 | 900 ± 400 |
| $[RGSGG]_5$ | 0.42 ± 0.06 | N/A |
| $[RGPGG]_5$ | 0.19 ± 0.08 | N/A |
| $[RPRPP]_5$ | 1.3 ± 0.2 | N/A |
| $[RARAA]_5$ | 6 ± 1 | 50 ± 20 |
| $[RQRQQ]_5$ | 2.8 + 0.4 | 19 ± 7 |
| $[RLRLL]_5$ | N/A | N/A |
| RG1Y | 8 ± 1 | 90 ± 30 |
| RG3Y | 13 ± 3 | 230 ± 70 |

### Supplementary Figures

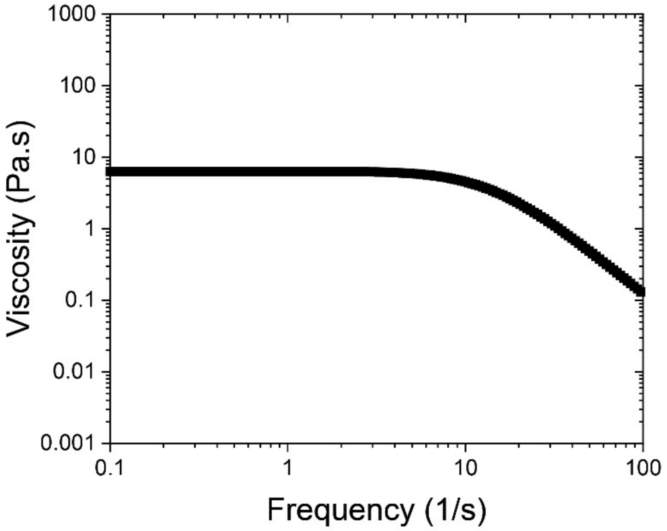


**Fig. S1**. The viscosity of $[RGRGG]_5$-rU40 condensates as a function of frequency as obtained from the pMOT experiments.

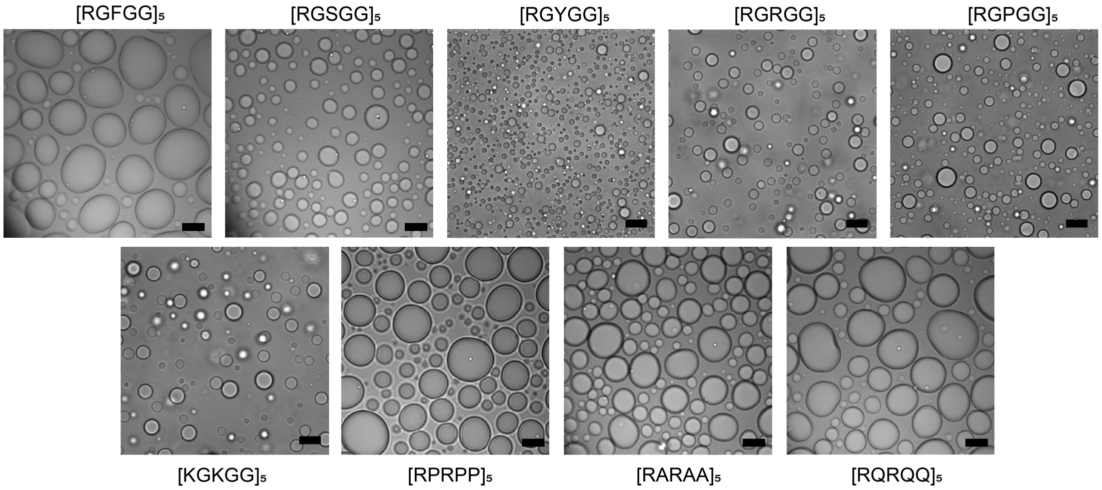


**Fig. S2**. Brightfield images showing spherical condensates formed by various peptides when mixed with rU40 RNA. Scale bars represent 10 µm.

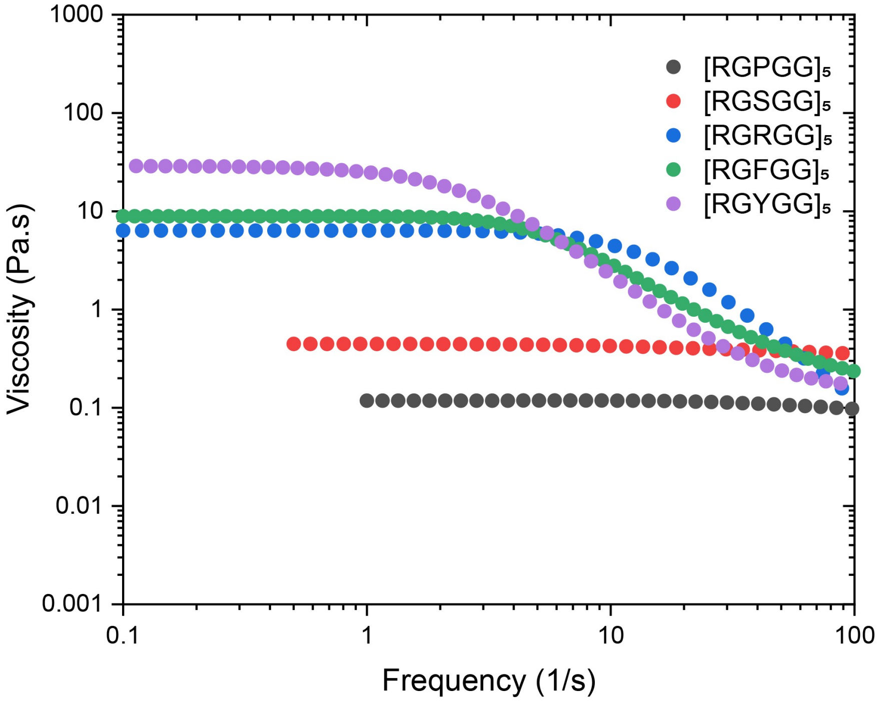


**Fig. S3**. The viscosity of RGXGG-rU40 condensates as a function of frequency as obtained from pMOT experiments. X is a variable amino acid that is set to P, S, R, F, or Y.

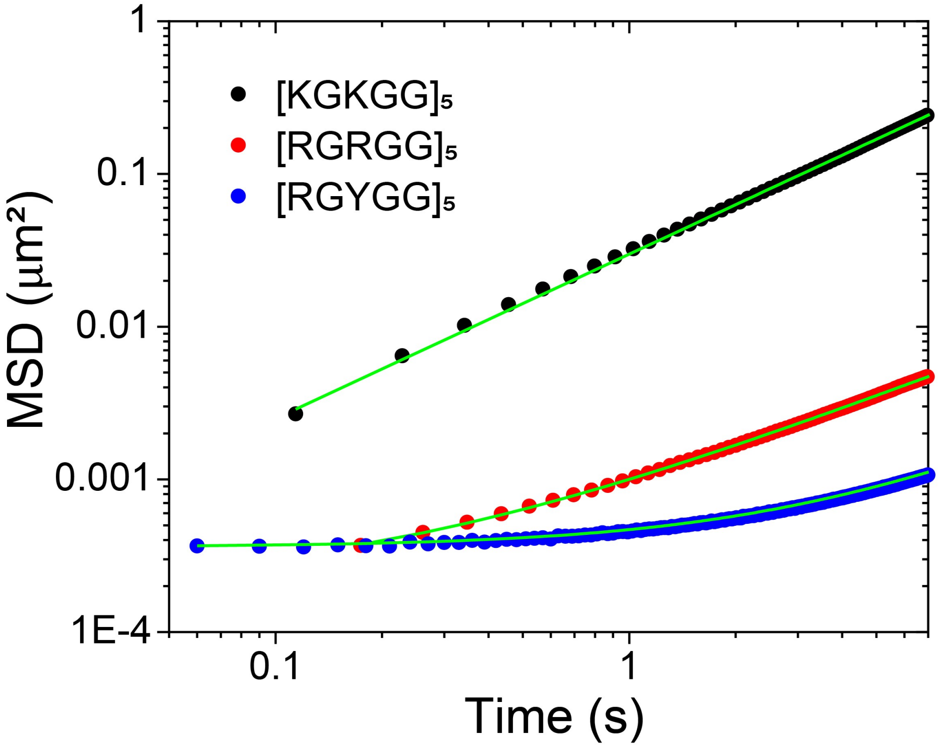


**Fig. S4**. The mean square displacement (MSD) of 200 nm polystyrene particles within condensates formed by rU40 and $[RGPGG]_5$ (black), $[RGRGG]_5$ (red), and $[RGYGG]_5$ (blue). Green lines are fits to the data using $MSD = 4Dt^{\alpha} + b$ for $[RGPGG]_5$ and $MSD = \Delta_0(1 + t/\tau)$ for $[RGYGG]_5$ and $[RGRGG]_5$ data[8].

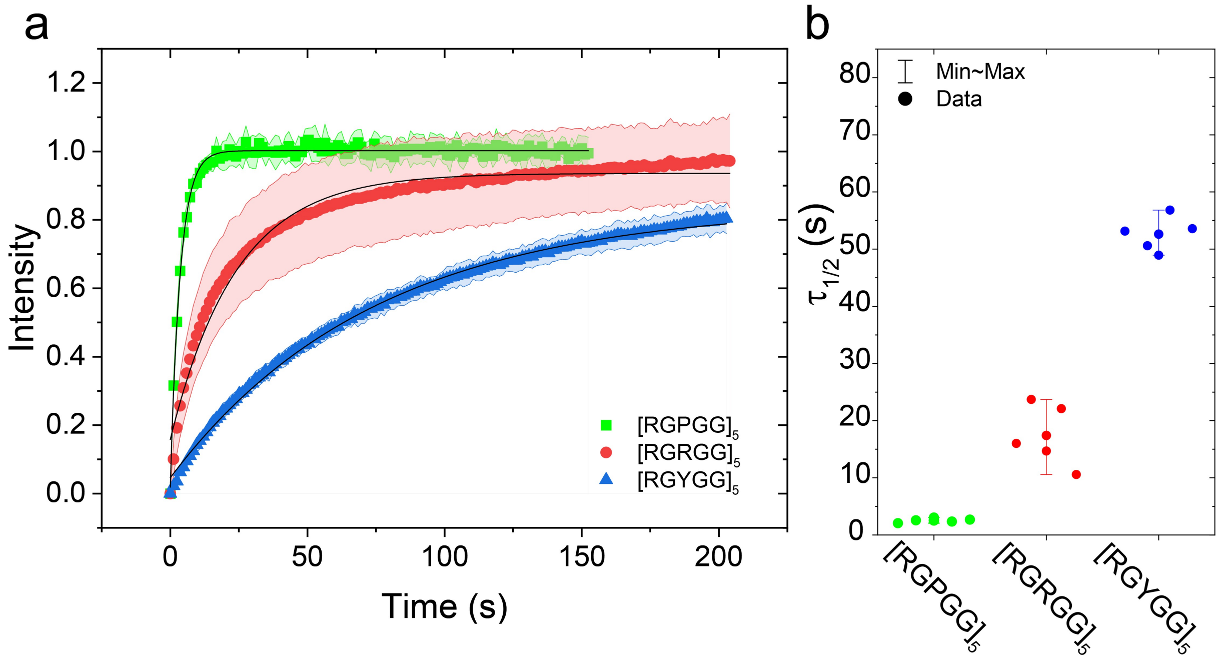


**Fig. S5**. **(a)** FRAP intensity-time traces of an RNA probe (rU10-FAM) in condensates formed by rU40 RNA and $[RGPGG]_5$ (green), $[RGRGG]_5$ (red), and $[RGYGG]_5$ (blue). Solid symbols are the average of five FRAP curves for each peptide-RNA combination. Error bars in the curves represent one s.d. obtained from the five curves for each peptide-RNA combination. Solid black lines are fits using $I(t) = A(1 - \exp(-t/\tau)$. **(b)** The recovery half-time of the same condensates as extracted from the data by fitting individual FRAP recovery traces. Error bars represent the range of the data.

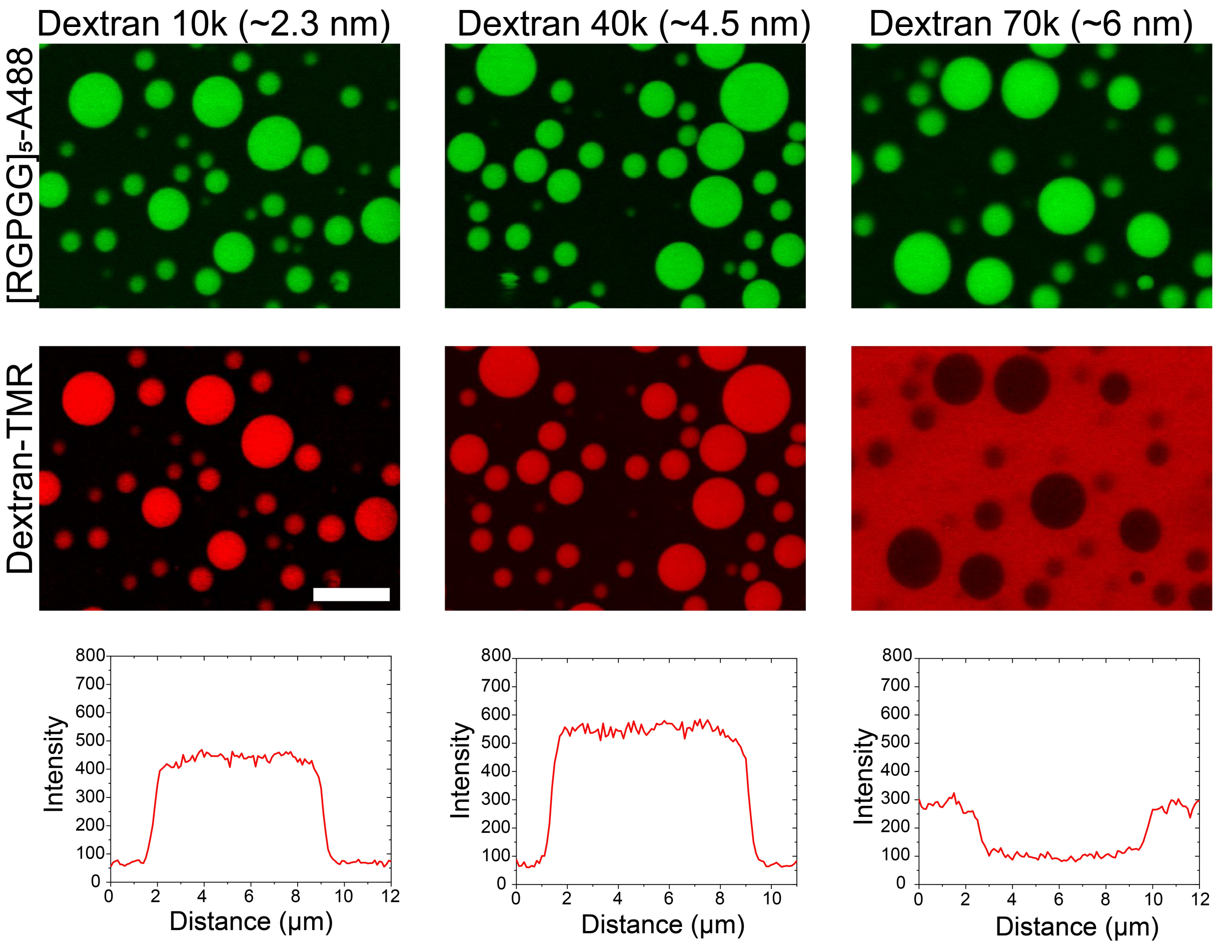


**Fig. S6.** Mesh size-determination experiment using TMR-labeled Dextran molecules with variable molecular weights for $[RGPGG]_5$-rU40 condensates. The upper panel shows the condensates as visualized by Alexa488-tagged $[RGPGG]_5$ (1% labeling ratio). The middle panel shows the partition behavior of Dextran molecules within peptide-RNA condensates. The scale bar represents 10 μm. The lower panel shows corresponding intensity profiles for Dextran molecules. The numbers in brackets indicate the estimated hydrodynamic radius of the Dextran molecules in aqueous solutions[10].

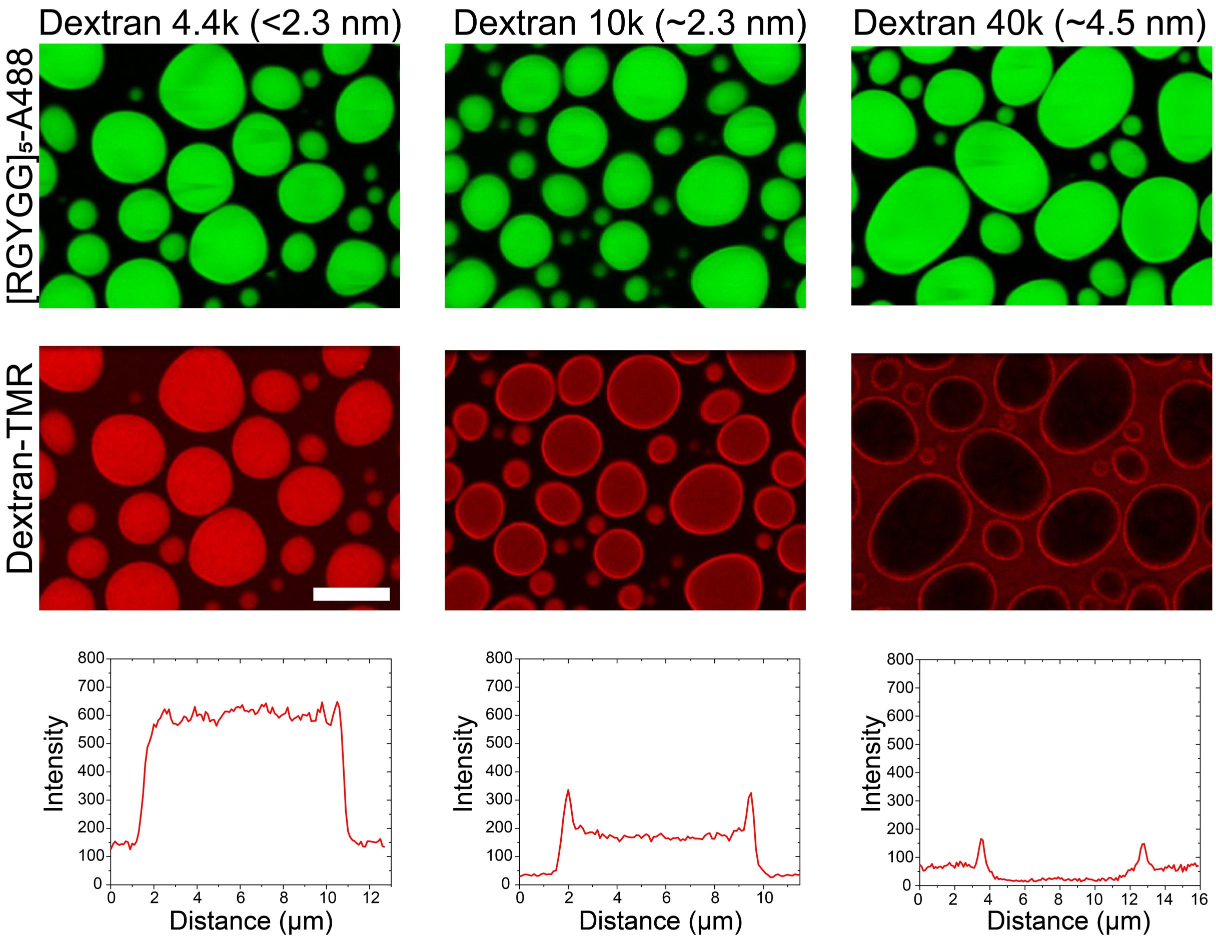


**Fig. S7.** Mesh size-determination experiment using TMR-labeled Dextran molecules with variable molecular weights for $[RGYGG]_5$-rU40 condensates. The upper panel shows the condensates as visualized by Alexa488-tagged $[RGYGG]_5$ (1% labeling ratio). The middle panel shows the partition behavior of Dextran molecules within peptide-RNA condensates. The scale bar represents 10 μm. The lower panel shows corresponding intensity profiles for Dextran molecules. The numbers in brackets indicate the estimated hydrodynamic radius of the Dextran molecules in aqueous solutions[10].

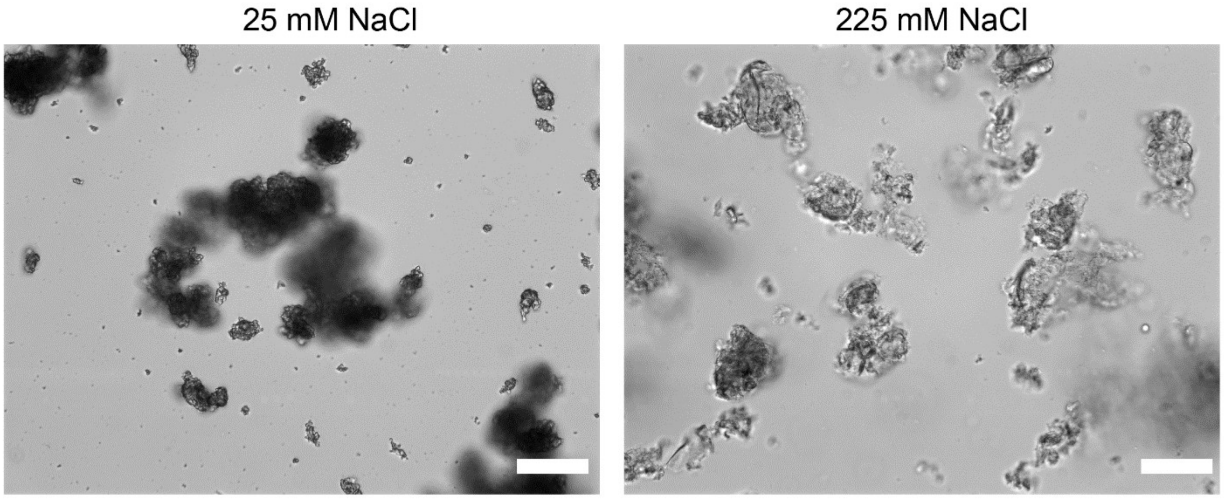


**Fig. S8**. Bright-field images of arrested condensate networks formed by [RLRLL]$_5$ and rU40 RNA at two different salt conditions in 25 mM Tris-HCl buffer (pH 7.5) and 20 mM DTT. Scale bars represent 20 μm.

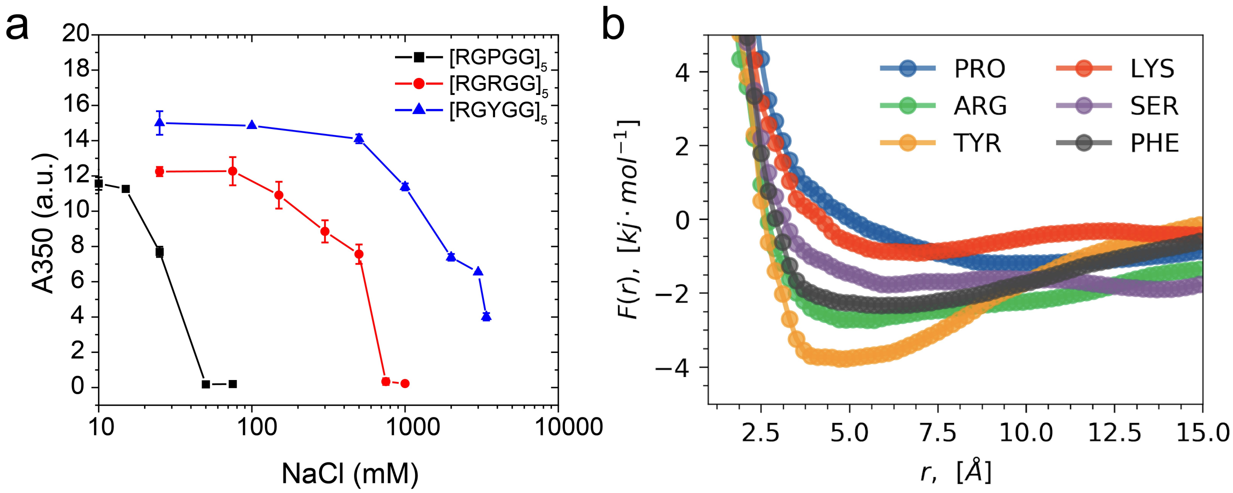


**Fig. S9. (a)** Solution turbidity at 350 nm wavelength (A350) for peptide-rU40 condensate as a function of salt concentration in 25 mM Tris-HCl buffer (pH 7.5). Data are shown for $[RGPGG]_5$, $[RGRGG]_5$, and $[RGYGG]_5$ condensates with rU40 RNA. **(b)** Free energy profiles of residue-rU attraction from model (GXG)-rU3 all-atom constant temperature simulations. X is set to Pro, Arg, Tyr, Lys, Ser, and Phe.

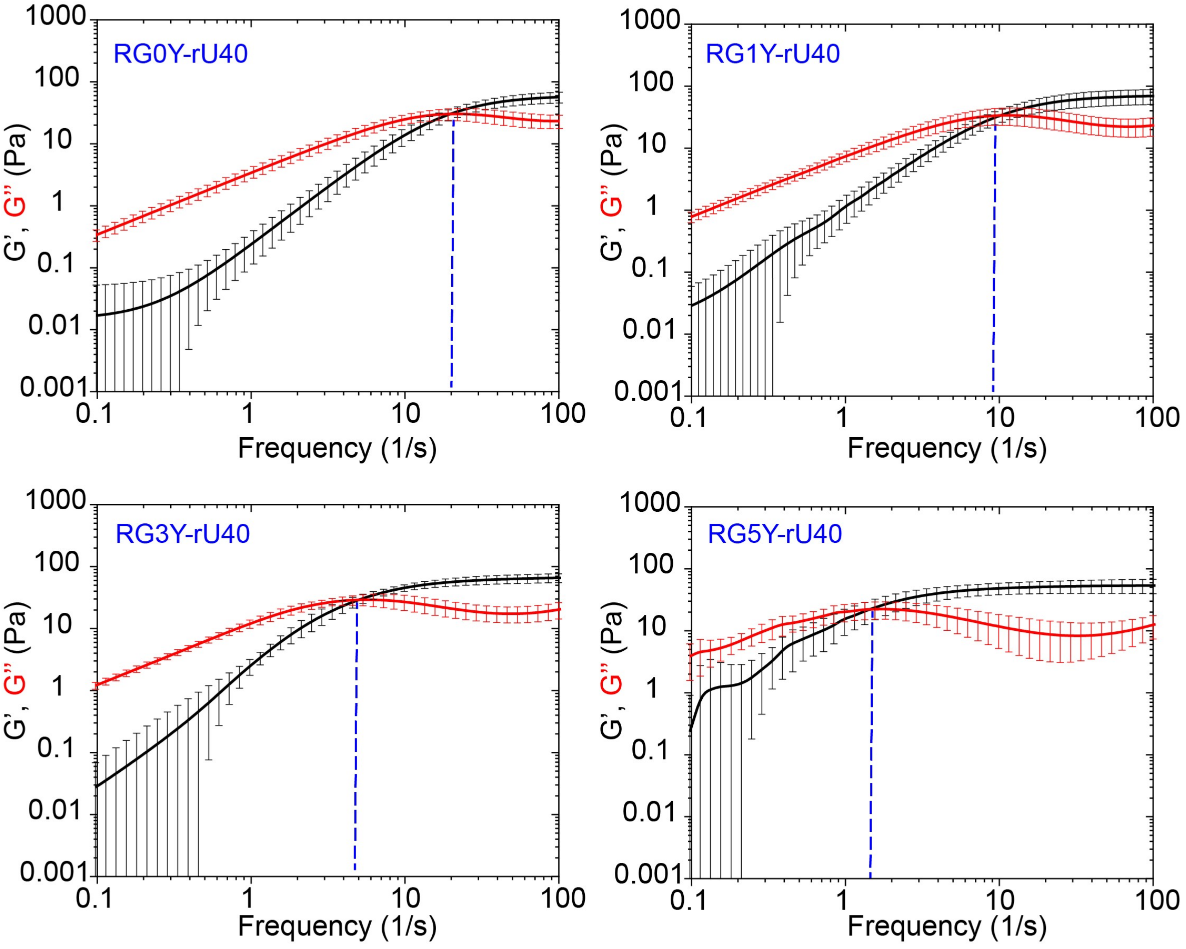


**Fig. S10**. Frequency-dependent viscoelastic moduli as obtained from pMOT experiments for condensates formed by RG$n$Y and RNA (see Table S1 for peptide sequences). Here $n$ is the number of R-to-Y mutations in the 2nd arginine of RGRGG-repeat peptide. $[RG\underline{R}GG]_5$ corresponds to RG0Y and RG5Y corresponds to $[RG\underline{Y}GG]_5$. The blue dashed lines indicate the crossover frequency. Error bars represent ±1 s.d.

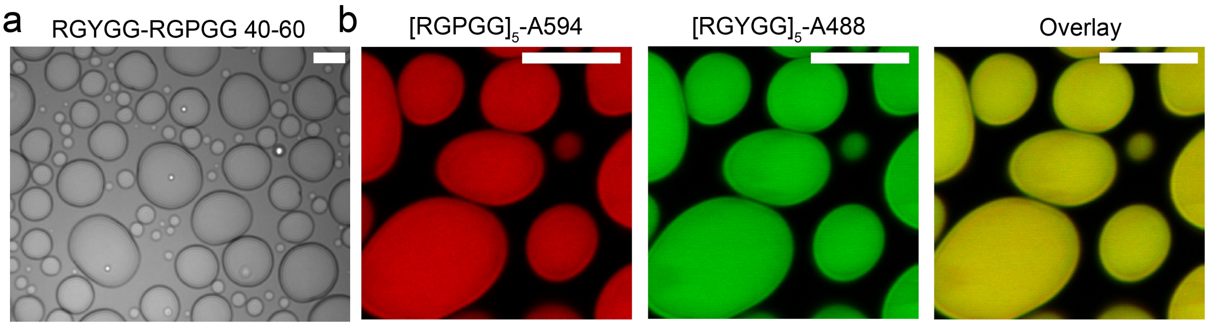


**Fig. S11**. **(a)** Brightfield image of condensates formed by $[RGYGG]_5$, $[RGPGG]_5$, and rU40 RNA. The total peptide concentration is 5 mg/ml (with 40% $[RGYGG]_5$ and 60% $[RGYGG]_5$). The RNA concentration is 2.5 mg/ml. The image shows that these droplets are homogenous and well-mixed. The sample also contained 1 μm beads for pMOT experiments. **(b)** Multi-color fluorescence images of condensates formed by mixtures of $[RGYGG]_5$ and $[RGPGG]_5$ in the presence of rU40 RNA. In this sample, the RGPGG-to-RGYGG ratio is 50:50. ~ 1% of Alexa-fluorophore-labeled peptides are used for imaging, as indicated. The scale bar is 10 μm.

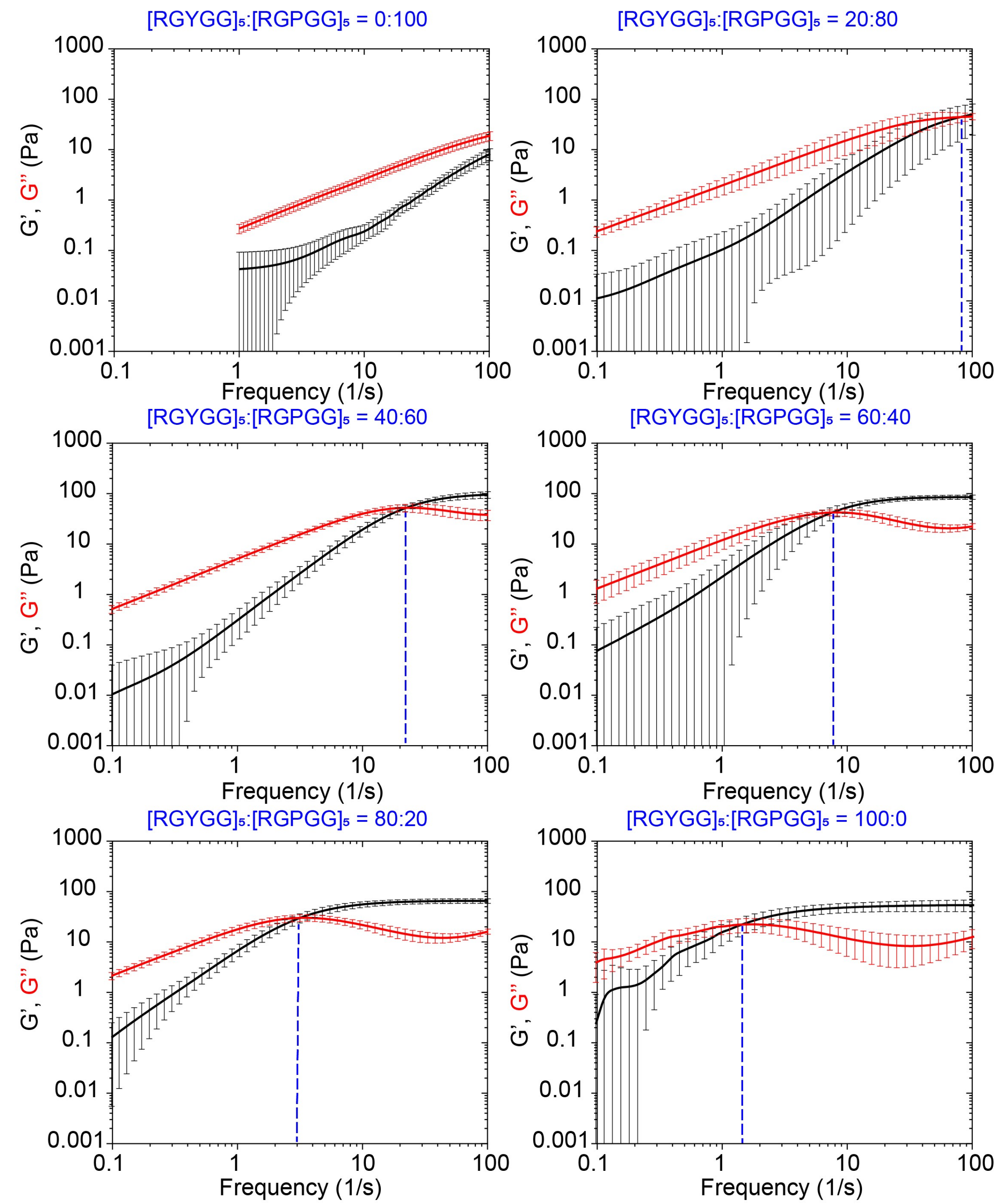


**Fig. S12**. Frequency-dependent viscoelastic moduli as obtained from pMOT experiments for condensates formed in a ternary mixture of $[RGYGG]_5$, $[RGPGG]_5$, and rU40. In these mixtures, the total peptide concentration is fixed at 5.0 mg/ml, the total rU40 concentration is 2.5 mg/ml. The molar fractions of $[RGYGG]_5$ and $[RGPGG]_5$ are varied while keeping the overall peptide concentration fixed to illustrate the composition-dependent viscoelastic modulation of these condensates. The blue dashed lines indicate the crossover frequency. Error bars represent ±1 s.d.

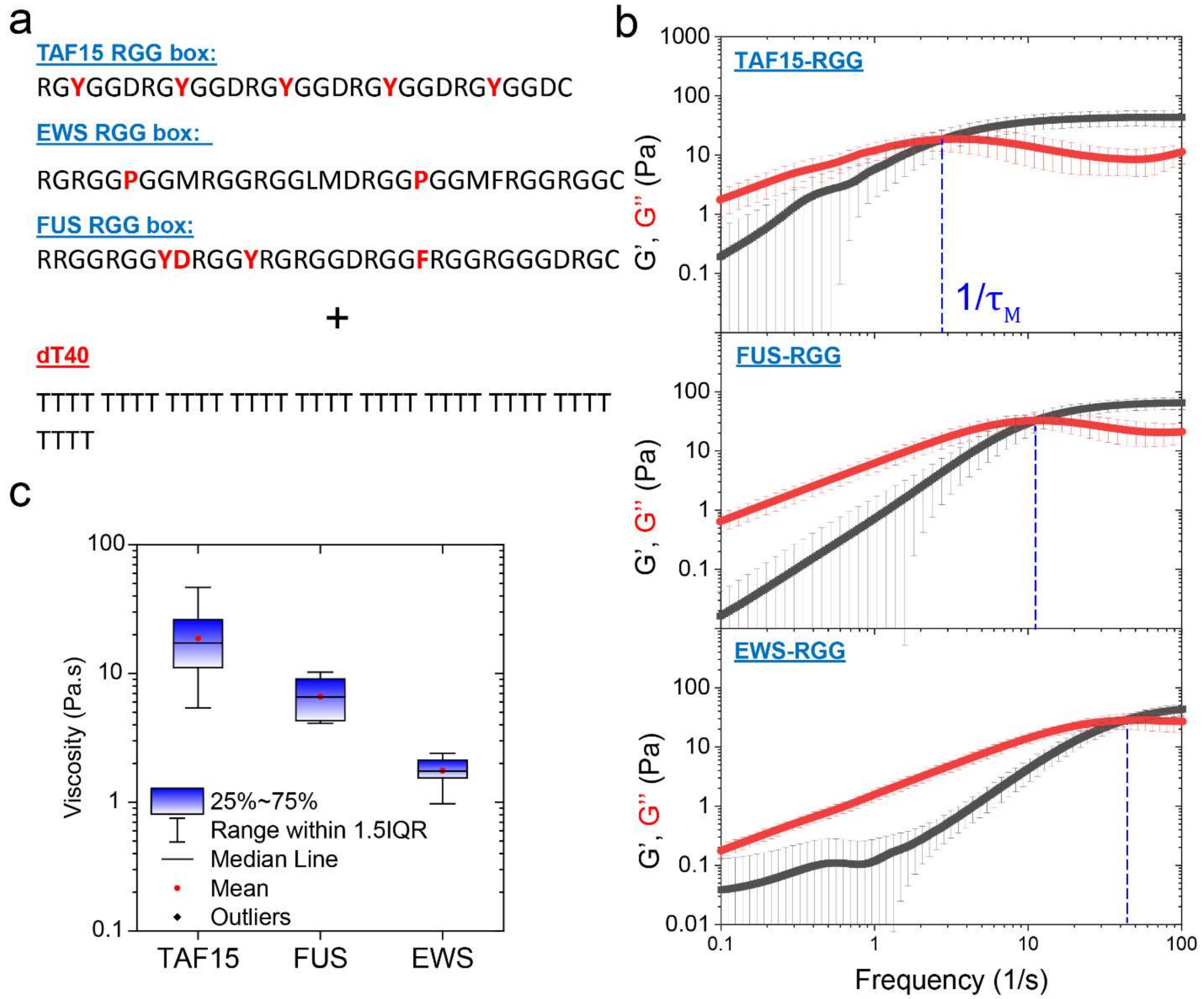


**Fig. S13.** pMOT measurements on the RGG boxes of FET proteins showing distinct viscoelastic properties. **(a)** Sequences of the RGG boxes derived from FUS, EWS, and TAF15. **(b)** Viscoelastic moduli of condensates formed by TAF15-RGG, FUS-RGG, or EWS-RGG with dT40. The blue dashed line indicates the inverse terminal relaxation time. Error bars represent ±1 s.d. **(c)** Terminal viscosity of condensates formed by TAF15-RGG, FUS-RGG, or EWS-RGG with dT40.

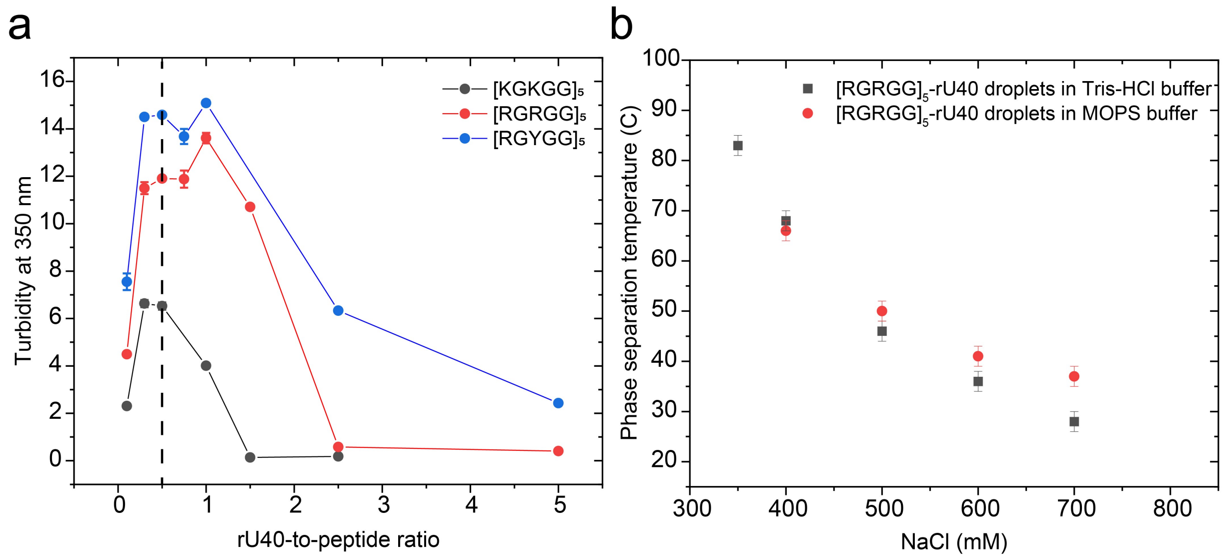


**Fig. S14**. **(a)** Turbidity of peptide-RNA mixtures as a function of RNA (rU40)-to-peptide weight ratio. The data are shown for $[KGKGG]_5$, $[RGRGG]_5$, and $[RGYGG]_5$ mixtures with rU40 RNA. The vertical dashed line represents the chosen ratio for the experiments in this study (RNA-to-peptide ratio of 0.5). **(b)** Comparison of phase separation temperature ($T_{ph}$) for $[RGRGG]_5$-rU40 condensates mixed in a buffer containing 25 mM Tris-HCl (pH 7.4) with variable salt and a buffer containing 25 mM MOPS (pH 7.4) and variable salt concentration. Both buffers also contain 20 mM DTT. This comparison is shown to assess the effect of temperature on the pH of the Tris buffer that might alter the phase separation behavior. The Tris buffer shows a similar trend of phase separation temperatures as compared to the MOPS buffer which has lesser pH sensitivity to temperature[15].

### Supplementary Note 1

*Data analysis:*

The analysis of pMOT experiments is summarized in a flow chart and shown in Figure S15. The output of a pMOT experiment is a trajectory of the bead (Fig. S15a) constrained by the harmonic potential of an optical trap within a peptide-RNA condensate. Each trajectory was analyzed in two steps: (i) calibration of the optical tweezer, and (ii) extraction of the condensate viscoelastic moduli. Both steps are done on the same trajectory.

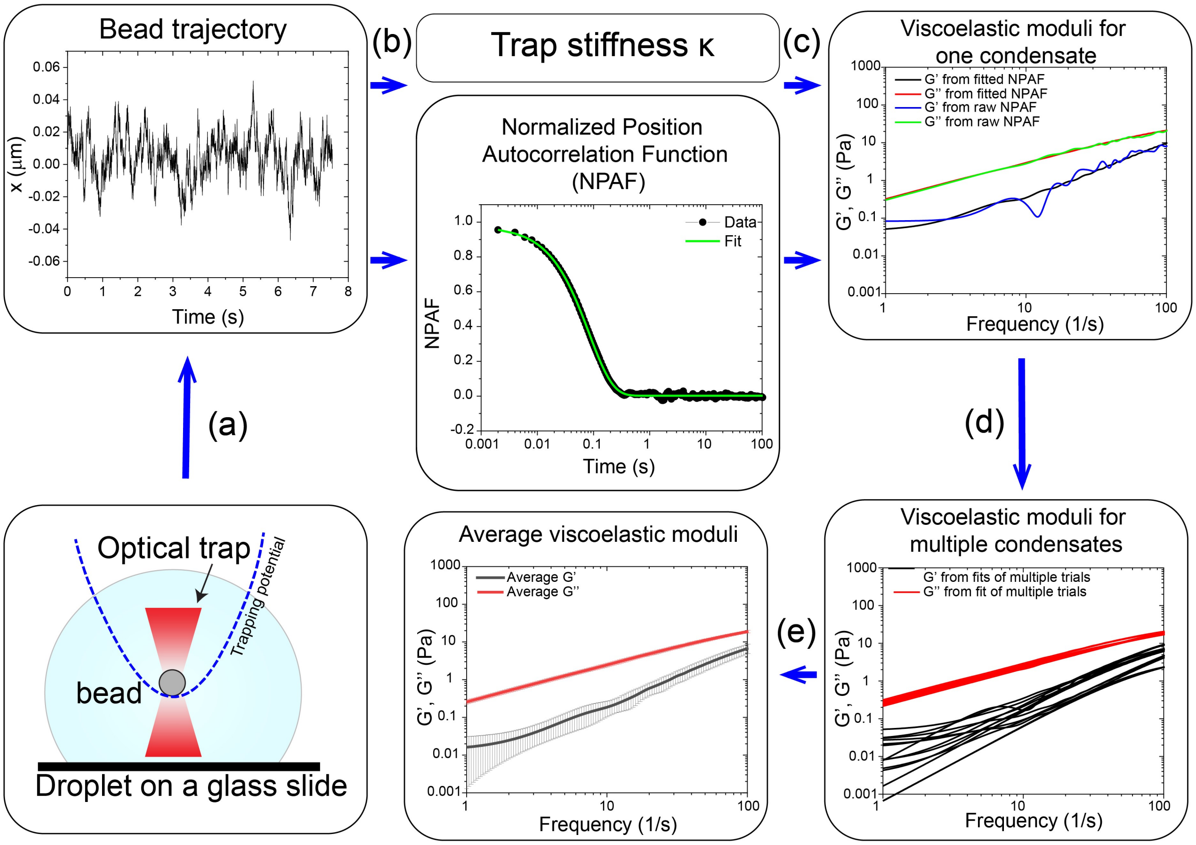


**Fig. S15**. Flow chart showing pMOT data analysis procedure to obtain the viscoelastic moduli of phase-separated condensates. **(a-e)** various steps of the procedure are described in the text of SI Note-1.

When a bead is diffusing within the harmonic potential of an optical trap, the motion of the bead is primarily mediated by the thermal fluctuations of the medium. According to the equipartition theorem, the variance of the position of the bead in one direction (X or Y) is given by the following equation[3]

$$\frac{1}{2}\kappa\langle x^2\rangle = \frac{1}{2}k_B T \quad (1)$$

Where $\kappa$ is the trap stiffness, $k_B$ is the Boltzmann constant, and $T$ is the temperature. For each recorded trajectory, the variance in both the X and Y coordinates was calculated from the trajectory. The centroid position accuracy was measured to be 1.5 nm (Fig. S16). The temperature of the stage was manually measured using a thermocouple and found to be 24 ± 0.5 °C. We ensured only using trajectories for which the variance in the X and Y directions were identical, indicating that the trapping potential is circular in the XY plane and therefore, the estimation of the trap stiffness from the harmonic form is valid. The values for the trap stiffness in the X and Y directions were compared and found to be consistent with each other for each measurement. This calibration method is one of the few methods that do not require a priori knowledge of the viscosity of the medium[3].

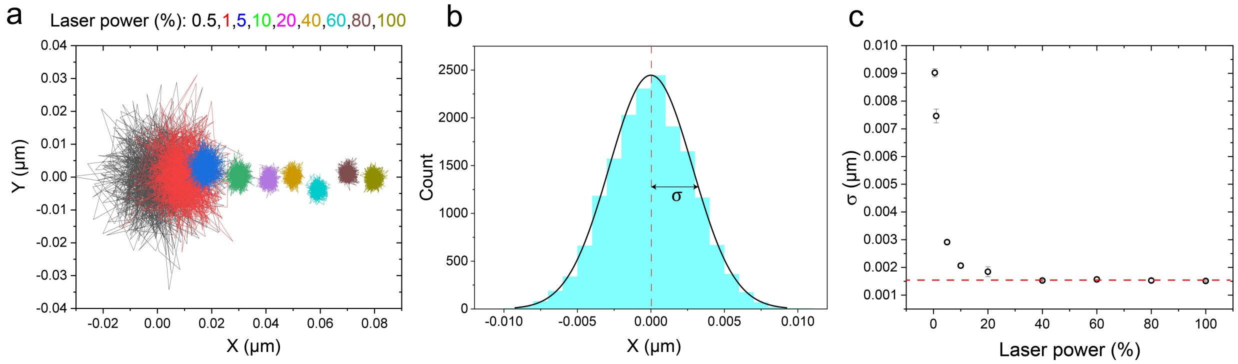


**Fig. S16. (a)** Trajectories of a bead trapped in water at different laser trapping powers (100% laser is ~1 mW). **(b)** Displacement distribution of the bead from the center of the trap. Data is shown for trapping power of 5 %. σ is the standard deviation of the fluctuations. **(c)** The standard deviation ($\sigma = \sqrt{\text{variance}}$) of the trapped bead position fluctuations under different laser trapping power values. The values of σ saturate at 1.5 nm, which is taken as the centroid positioning accuracy.

For a particle undergoing Brownian motion in a fluid and constrained by an optical trap, the particle's motion can be described by a generalized Langevin equation similar to the one described by Mason and Wietz[6] with an added term for the optical trap[3]

$$m\vec{a}(t) = \overrightarrow{f_R}(t) - \int_0^t \zeta(t-\tau)\vec{v}(\tau)\,d\tau - \kappa\vec{r}(t) \quad (2)$$

Where $m$ is the mass of the particle and $\vec{a}$ is the acceleration. The first term $\overrightarrow{f_R}(t)$ represents a white noise function to model the thermal fluctuations. The second term involves a memory function $\zeta(t-\tau)$ that represents damping forces exerted by the fluid on the particle [$\vec{v}(\tau)$ is the particle velocity], and the third term is the optical trap force. In one dimension, equation (2) can be rewritten as a differential equation in terms of the position of the particle[3-5]

$$m\ddot{r}(t) = f_R(t) - \int_0^t \zeta(t-\tau)\dot{r}(\tau)\,d\tau - \kappa r(t) \quad (3)$$

Where the dot accent represents a time derivative. To solve this equation, the method of Laplace transforms is used. The solution in the Laplace space is given by[3-5]

$$\tilde{\zeta}(s) = \kappa\frac{\tilde{A}(s)}{1-s\tilde{A}(s)} - ms \quad (4)$$

Where $A(s)$ is the normalized position autocorrelation function in the Laplace space. At the same time, the memory function was evaluated by Mason and Weitz in relation to the viscosity of the medium under equilibrium conditions as[6]

$$\tilde{\zeta}(s) = 6\pi a\tilde{\eta}(s) \tag{5}$$

Hence, combining the two equations (4 and 5) gives a relation between the viscosity and the autocorrelation function in the Laplace space[3]

$$\tilde{\eta}(s) = \frac{\kappa}{6\pi}\left(\frac{\tilde{A}(s)}{1-s\tilde{A}(s)} - \frac{ms}{\kappa}\right) \tag{6}$$

The viscosity in the Laplace space can be transformed into the complex modulus in the Fourier space using[3]

$$G^*(\omega) = s\tilde{\eta}(s)|_{s=i\omega} \tag{7}$$

Thus, the final expression for the complex modulus in terms of the particle normalized position autocorrelation function is expressed as[2,3,5]

$$G^*(\omega) = \frac{\kappa}{6\pi a}\left(\frac{i\omega\hat{A}(\omega)}{1-i\omega\hat{A}(\omega)} + \frac{m\omega^2}{\kappa}\right) \tag{8}$$

Where $\kappa$ is the stiffness of the trap and $a$ is the particle radius. The second term in the bracket is negligible except for very high frequencies $\omega \sim 1\, MHz$. Therefore, we used this equation (neglecting the second term) to obtain the complex shear modulus of the material[2,3,5]

$$G^*(\omega) = G'(\omega) + i\, G''(\omega) = \frac{\kappa}{6\pi a}\left(\frac{i\omega\hat{A}(\omega)}{1-i\omega\hat{A}(\omega)}\right) \tag{9}$$

Equation (9) was implemented in a custom-built python script and used to extract the elastic modulus $G'(\omega)$ and the viscous modulus $G''(\omega)$ of the condensate. The frequency-dependent viscosity $\eta(\omega)$ was calculated using[5]

$$\eta(\omega) = \frac{G''(\omega)}{\omega} \tag{10}$$

The zero-shear viscosity was obtained from the limit of $\eta(\omega)$ at low frequencies. We calculate the autocorrelation function $A(\tau)$ from a trajectory of a bead within a condensate in one coordinate (X or Y, Fig. S15b). This was done using the *multipletau.autocorrelate()* function from the multipletau python library[16] (v0.1.9). The autocorrelation function is fitted with a multi-exponential function to reduce the measurement noise (Fig. S15b and Figs. S22 and S23)

$$A(\tau) = A_1 \exp\left(-\frac{\tau}{b_1}\right)^{\alpha_1} + A_2 \exp\left(-\frac{\tau}{b_2}\right)^{\alpha_2} + A_3 \exp\left(-\frac{\tau}{b_3}\right)^{\alpha_3} + A_4 \tag{11}$$

Next, we perform a numerical Fourier transform on $A(\tau)$ to obtain $A(\omega)$. Following the procedure described by Evans et al.[17], for a function $g(t)$ with discrete points $(t_k, g_k)$ and boundary conditions $g(0) = 1$ and $\dot{g}(t) \to 0$ as $t \to \infty$,

$$-\omega^2\hat{g}(\omega) = i\omega g(0) + \frac{(1-e^{-i\omega t_1})(g_1 - g(0))}{t_1} + \dot{g}(\infty)e^{-i\omega t_N} + \sum_{k=2}^{N}\left(\frac{g_k - g_{k-1}}{t_k - t_{k-1}}\right)(e^{-i\omega t_{k-1}} - e^{-i\omega t_k}) \tag{12}$$

This equation was implemented to calculate $\hat{A}(\omega)$ from the fitted $A(\tau)$. Classical numerical Fourier transform methods require the function to have equidistant time points. However, this method of

Fourier transform does not have that requirement[17]. Finally, $A(\omega)$ was substituted in equation (9) to calculate the complex modulus $G^*(\omega) = G'(\omega) + iG''(\omega)$ (Fig. S15c and Figs. S24 and S25).

For each bead trajectory from each condensate, we extracted $G'(\omega), G''(\omega)$, and $\eta$ (Fig. S15d). The final viscoelastic moduli from multiple condensates were averaged for each frequency $\omega$. Error bars were estimated using the standard deviation (Fig. S15e). The presented values for the zero-shear viscosity are the average of at least 12 values extracted from individual condensates' moduli. The terminal relaxation time was obtained by averaging the inverse of the crossover frequency, which was calculated from G' and G'' frequency traces for individual condensates. An example of the condensate-to-condensate variation of the moduli is depicted in Figure S15d. To ensure the correctness of our calculations, we used this method to measure the viscosity of water found it to be 0.93 ± 0.03 mPa.s which is comparable with the reported value in the literature[18] (0.91 mPa.s, Fig. S17).

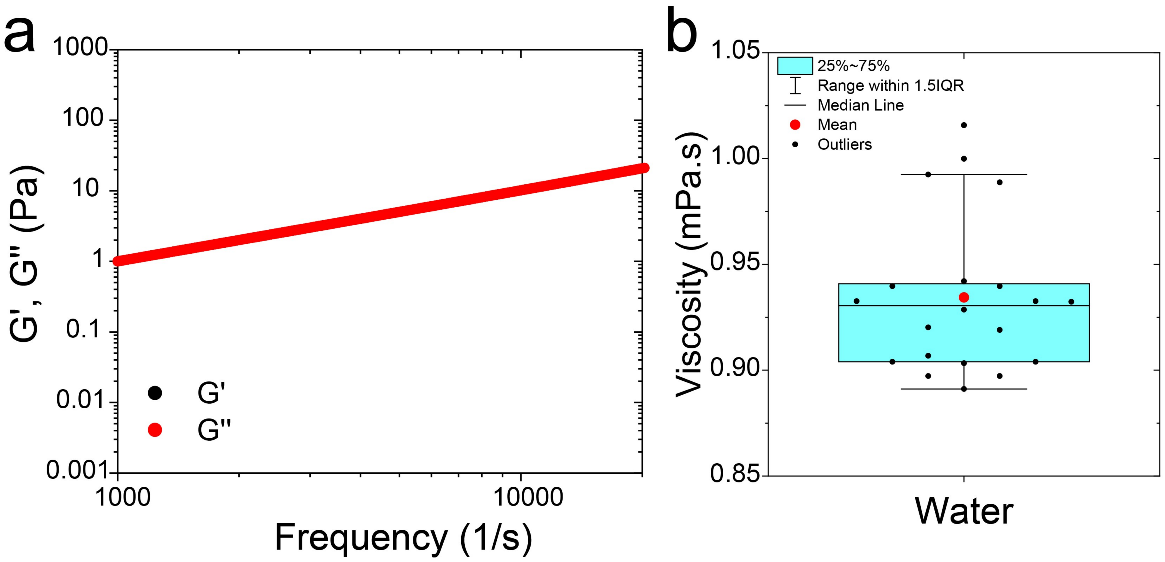


**Fig. S17**. **(a)** Viscoelastic moduli of water using the pMOT assay with 2 μm polystyrene beads. **(b)** The viscosity of water as measured from the pMOT assay (water viscosity is 0.91 mPa.s[18]).

***Bead condensate interactions***:

One of the assumptions of microrheology is that there are no strong interactions between the bead and condensate medium. The presence of such interactions alters the extracted rheological response of the material[19]. We have previously probed the effect of bead-condensate interactions on video particle tracking microrheology using $[RGRGG]_5$-dT40 condensates. In that study, we showed that the presence of the carboxylate beads did not alter the overall phase behavior of the peptide-nucleic acid system (Figure S3 in Alshareedah et al.[1]). This means that the beads are not affecting the interactions between the nucleic acids and the peptide significantly. To further confirm that, we performed a bead-halo assay on non-fluorescent carboxylate beads embedded within peptide-RNA condensates. The bead halo assay is used to probe intermolecular interactions by coating a bead with a substrate and adding the bait protein[20]. If the protein binds to the substrate it will coat the bead and the bead surface will have an intensity higher than the average bulk intensity[20] (Fig. S18a). We used carboxylate functionalized beads which partition positively in peptide-RNA condensates. As our baits, we used $[RGRGG]_5$-Alexa594 and U10-FAM RNA to see if the beads are interacting with the peptide or the RNA inside the condensates. Our experiments show that beads' intensity is less than or equal to the mean intensity of the probes inside the condensates, indicating no significant interactions or adsorption of the biopolymers on the bead surface (Fig. S18b). This result is also consistent with a recent study that showed that the MSD of PEGylated beads and carboxylate beads is identical within condensates formed by polyR10 ($[R]_{10}$) and UTP[21].

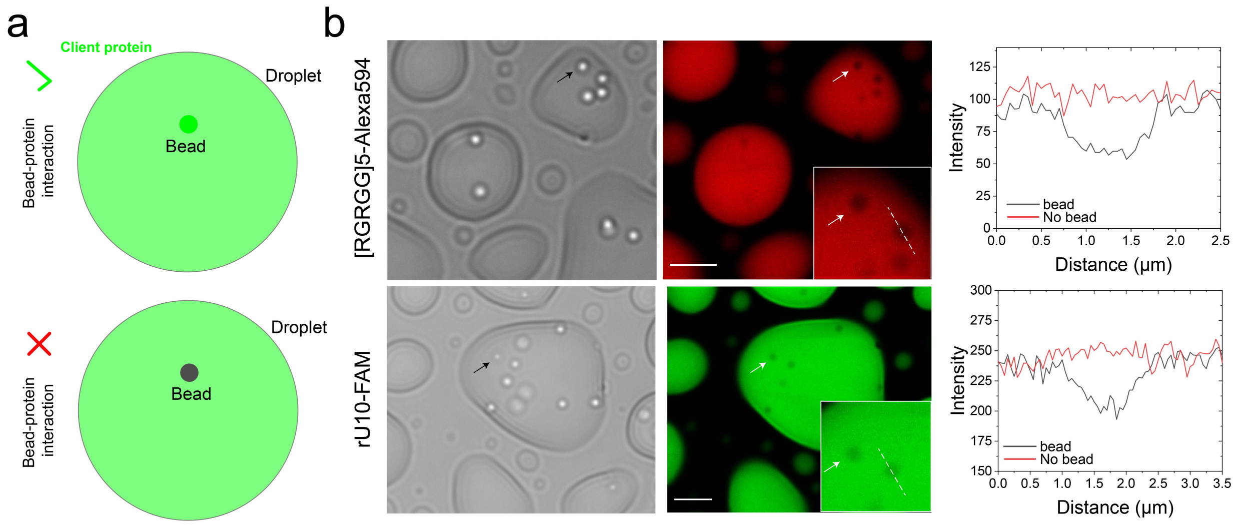


**Figure S18. (a)** Schematic illustration of the expected outcome of the bead-halo assay in case of the presence and absence of bead-client interactions. **(b)** Bright-field and fluorescent images of carboxylate beads within $[RGRGG]_5$-rU40 condensates and corresponding intensity profiles. (*Top panel*) The recruitment behavior of $[RGRGG]_5$-Alexa594 with the carboxylate beads within peptide-RNA condensates showing that the beads do not recruit or adsorb any peptide molecules on its surface. The intensity profile across the bead (black) shows a dip in intensity due to the absence of any $[RGRGG]_5$-Alexa594 molecules on its surface. The intensity profile across a bead-free region (red) of the droplets shows the average mean intensity. (*Bottom panel*) similar data but with U10-FAM RNA as the client in the bead-halo assay. The scale bars represent 10 µm. These results indicate that the carboxylate beads do not interact significantly with the condensate medium as they don't recruit or concentrate the two components forming the condensates: $[RGRGG]_5$ and RNA.

***Effect of solid and liquid interfaces:***

For accurate rheological analysis, the effects of nearby solid and liquid interfaces need to be quantified in order to ensure that laser refraction and interfacial fluctuations do not have significant effects on the measurements. First, we confirmed that the proximity of a liquid-liquid interface (~3 µm) does not significantly alter the viscoelastic moduli (Fig. S19), even though all of our experiments were performed when the bead is at the center of the condensate (distance from a liquid-liquid interface > 3 μm).

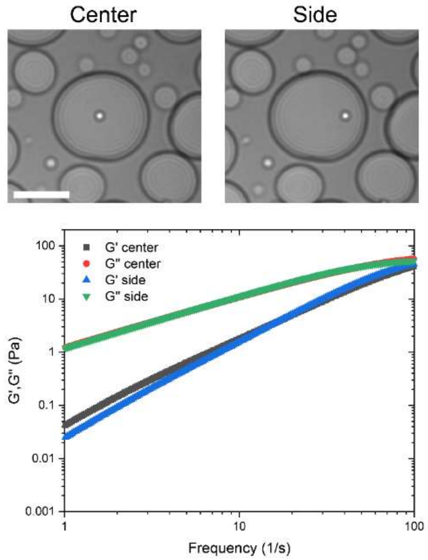


**Fig. S19**. A comparison between the frequency-dependent viscoelastic moduli (G' and G'') as obtained from pMOT experiments at the center of the condensate and at the side of the condensate (~ 3 μm from the condensate interface with the dilute phase). These condensates are formed by mixtures of $[RPRPP]_5$ peptide (5.0 mg/ml) and rU40 RNA (2.5 mg/ml). No significant effects of the proximity to the interface on the viscoelastic moduli were observed under these conditions. The scale bar represents 10 μm.

Next, we probed whether the proximity of the solid glass interface affects the optical trapping and the pMOT measurements due to the reflection and refraction of the trapping laser as well as bead-surface interactions. First, we measured the normalized position autocorrelation function (NPAF) for 1 μm bead trapped at a variable distance from the glass surface $h$ in water using identical trapping power (Fig. S20a). We found that there is insignificant variation in the NPAF curves as we change the distance from the surface from 3 μm to 10 μm. The same result was obtained from an identical experiment using a 35% aqueous solution of polyethylene glycol (Fig. S20b&c). This indicates that the trajectory of the bead in the X and Y direction 3 μm away from the surface is similar to that when the bead is 10 μm away from the surface.

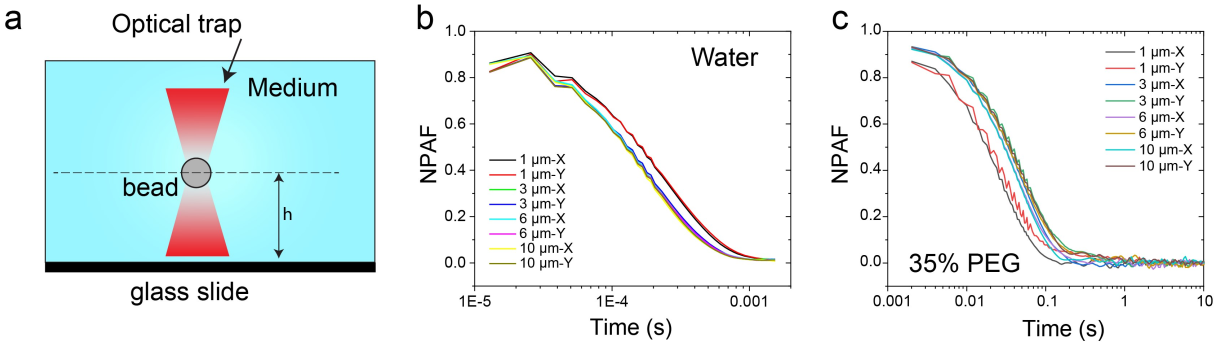

**Fig. S20.** (**a**) A scheme illustrating the trapping of a particle in a continuous medium at a variable distance $h$ from the glass surface. (**b**) Normalized position autocorrelation function (NPAF) of a trapped bead at a variable distance from the surface and at identical laser power within water. (**c**) Normalized position autocorrelation function (NPAF) of a trapped bead at a variable distance from the surface and at identical laser power within a solution of 35 % wt/vol PEG8000.

Next, we trapped a 1 μm bead within a peptide-RNA condensate ($[KGKGG]_5$-rU40) at different distances from the surface $h$ and measured the bead trajectory (Fig. S21a). We find that similar to the case of a continuous medium (Fig. S20), the NPAF curves were similar for the bead at all the tested surface to bead separations (1μm - 10μm; Fig. S21b). Additionally, the displacement probability in the X and Y direction was identical across all the tested surface-to-bead separations (Fig. S21c). This indicates that doing pMOT experiments under these conditions is not affected by the proximity of the glass coverslip surface nor by the liquid-liquid interface (Fig. S19). The Gaussian shape of the displacement probability also indicates that the trapping potential is harmonic[22]. Increasing the laser trapping power led to progressively narrower displacement probability distributions of the trapped bead within the peptide-RNA condensate (Fig. S21d). Furthermore, the relation between the measured trap stiffness and the laser power was found to be linear inside the condensate (Fig. S21e), which is expected for a harmonic optical trap[23]. Collectively, these data confirm the integrity of the optical trap when present within a peptide-RNA condensate and the validity of the harmonic potential approximation that facilitates our passive microrheology measurements.

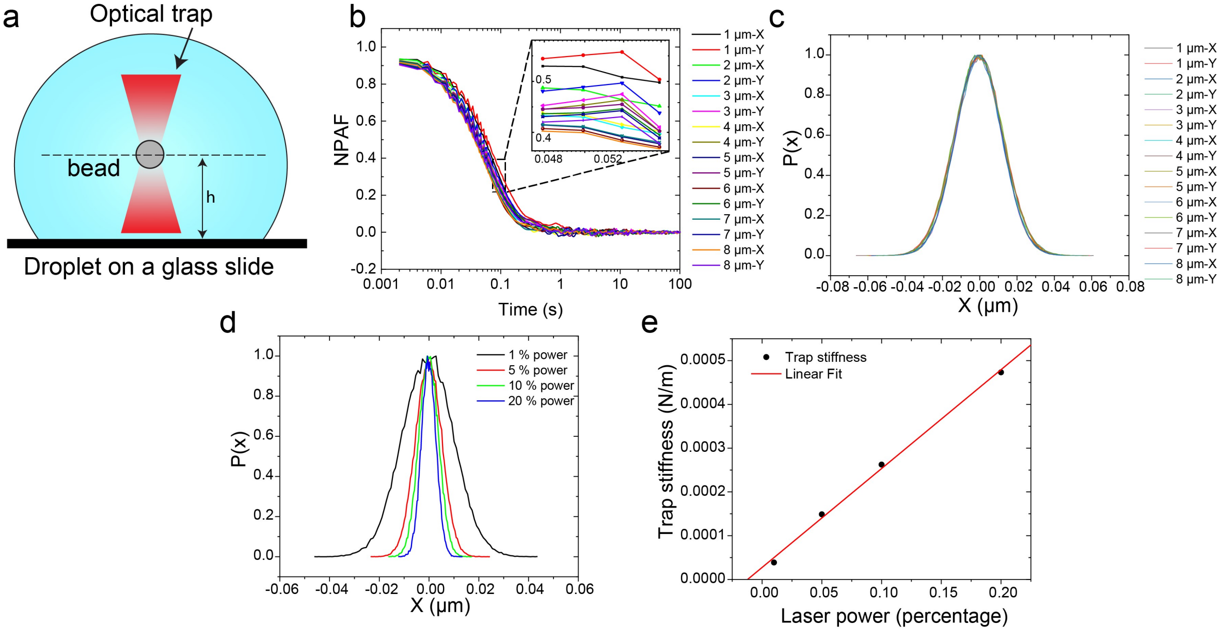


**Fig. S21.** Control experiments probing the feasibility of pMOT experiments within $[KGKGG]_5$-rU40 condensates. **(a)** Scheme illustrating the optical trapping of a polystyrene microsphere within peptide-RNA droplet at distance h from the glass slide surface. **(b)** Normalized position autocorrelation function (NPAF) of a trapped microsphere within the condensate at variable distances from the glass slide surface and identical trapping power. **(c)** Probability distribution of the trapped particle displacements from the center of the optical trap at variable distances from the glass slide surfaces. **(d)** The probability distribution of particle displacements from the center of the optical trap as a function of increasing trapping power and trapping stiffness within peptide-RNA condensate. **(e)** Trap stiffness scales linearly with laser power inside peptide-RNA condensates.

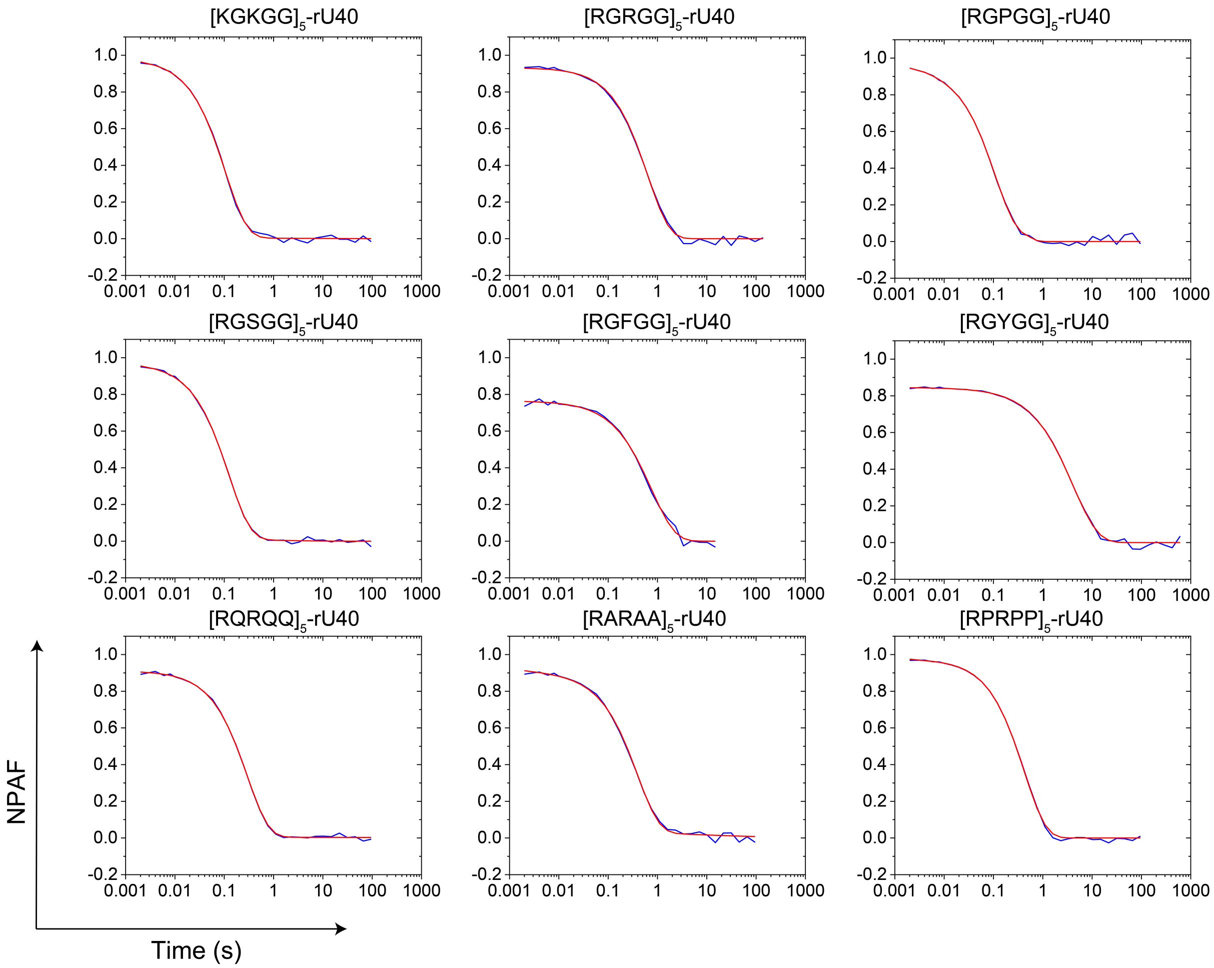


**Fig. S22.** Representative plots of the calculated normalized position autocorrelation function (NPAF) and the multi-exponential fit (Equation-11) for various peptide-RNA condensate systems that are shown in Figs. 1-3 in the main text.

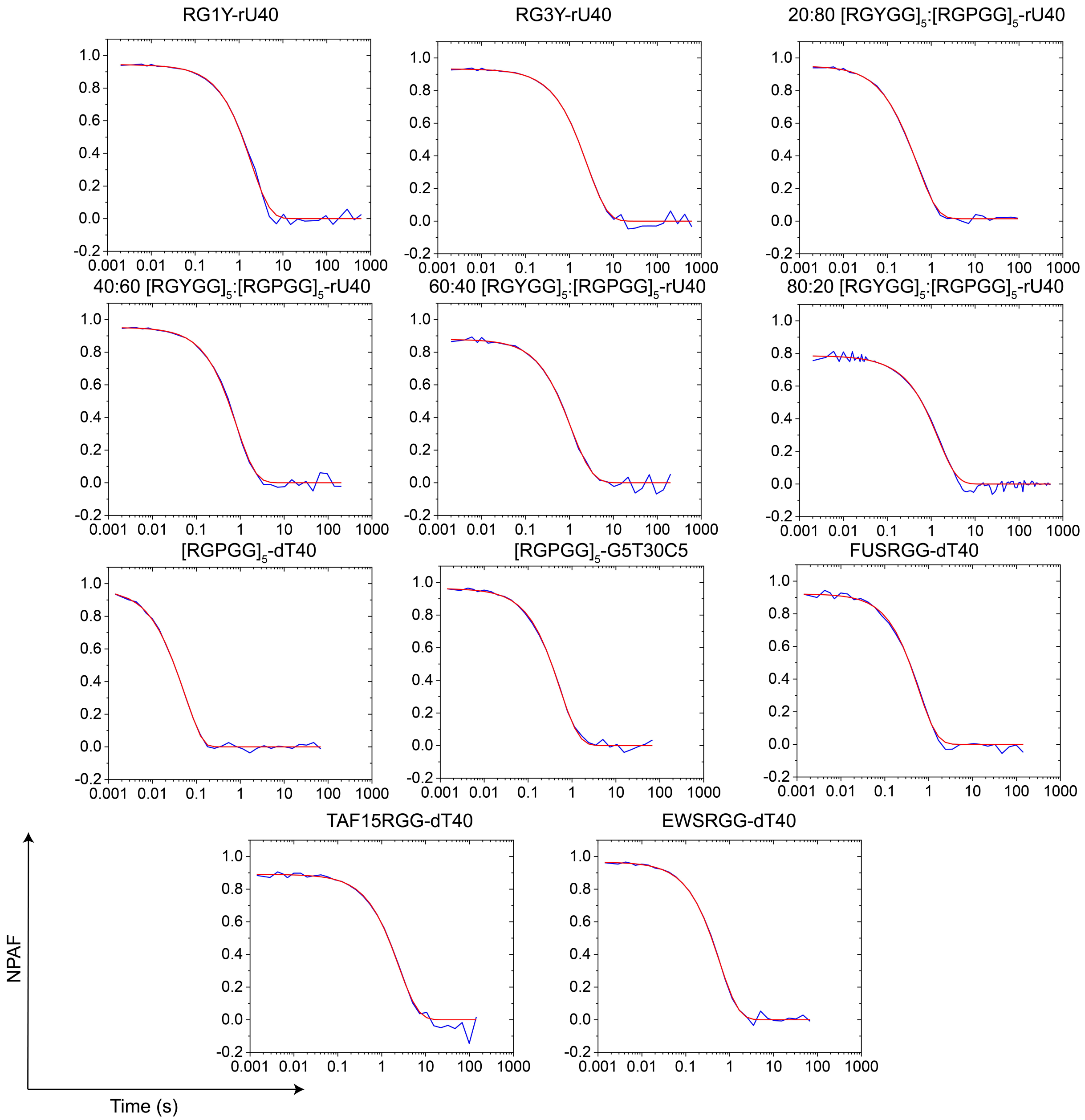


**Fig. S23.** Representative plots of the calculated normalized position autocorrelation function (NPAF) and the multi-exponential fit (Equation 11) for various peptide-NA condensate systems that are shown in Figs. 4-5 in the main text and corresponding SI figures.

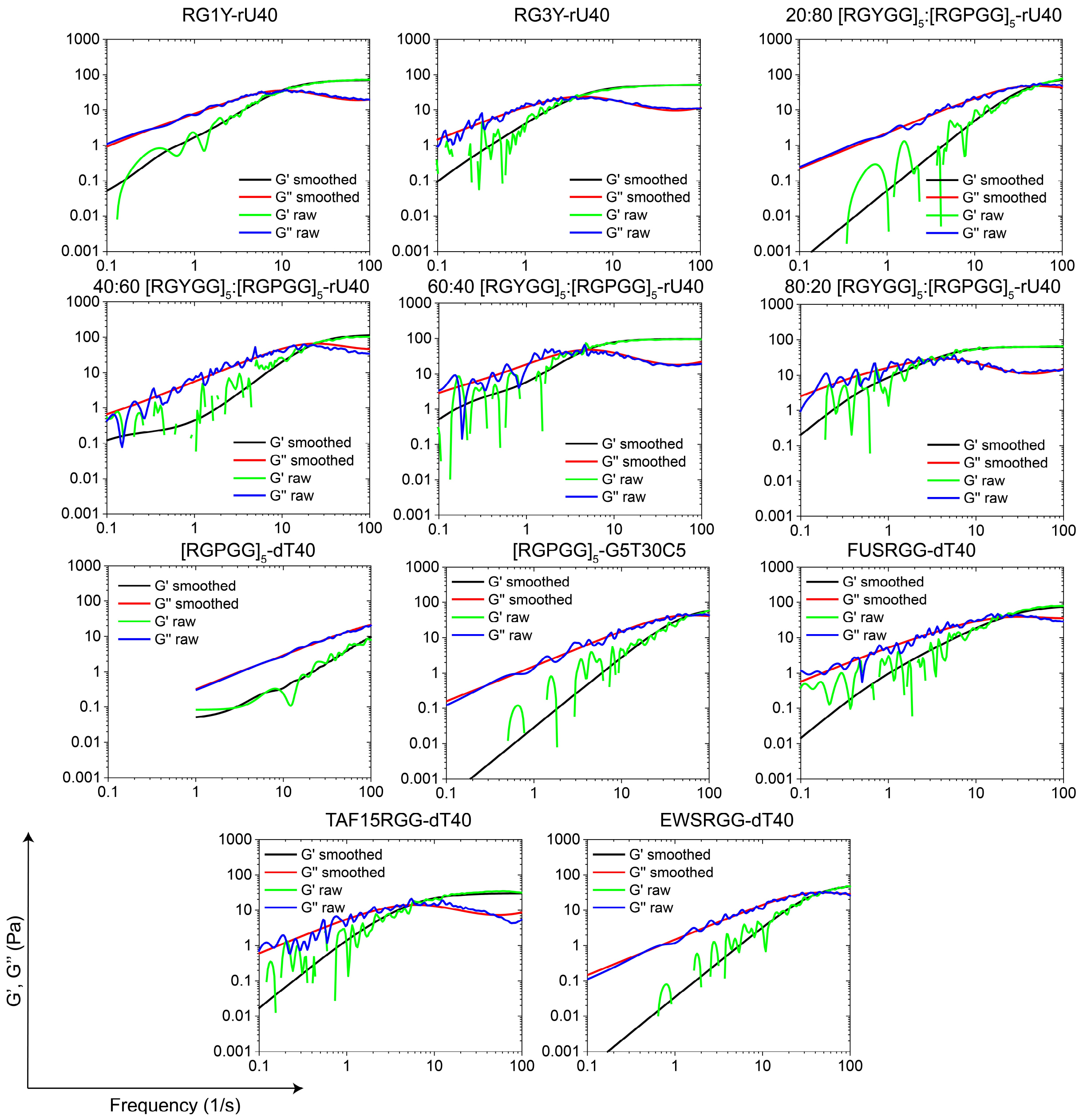


**Fig. S24.** Example plots of the viscoelastic moduli calculated from raw (blue and green) and fitted (black and red) autocorrelation functions (NPAF) of various peptide-NA condensate systems shown in Figs. 4-5 in the main text and corresponding supplementary figures.

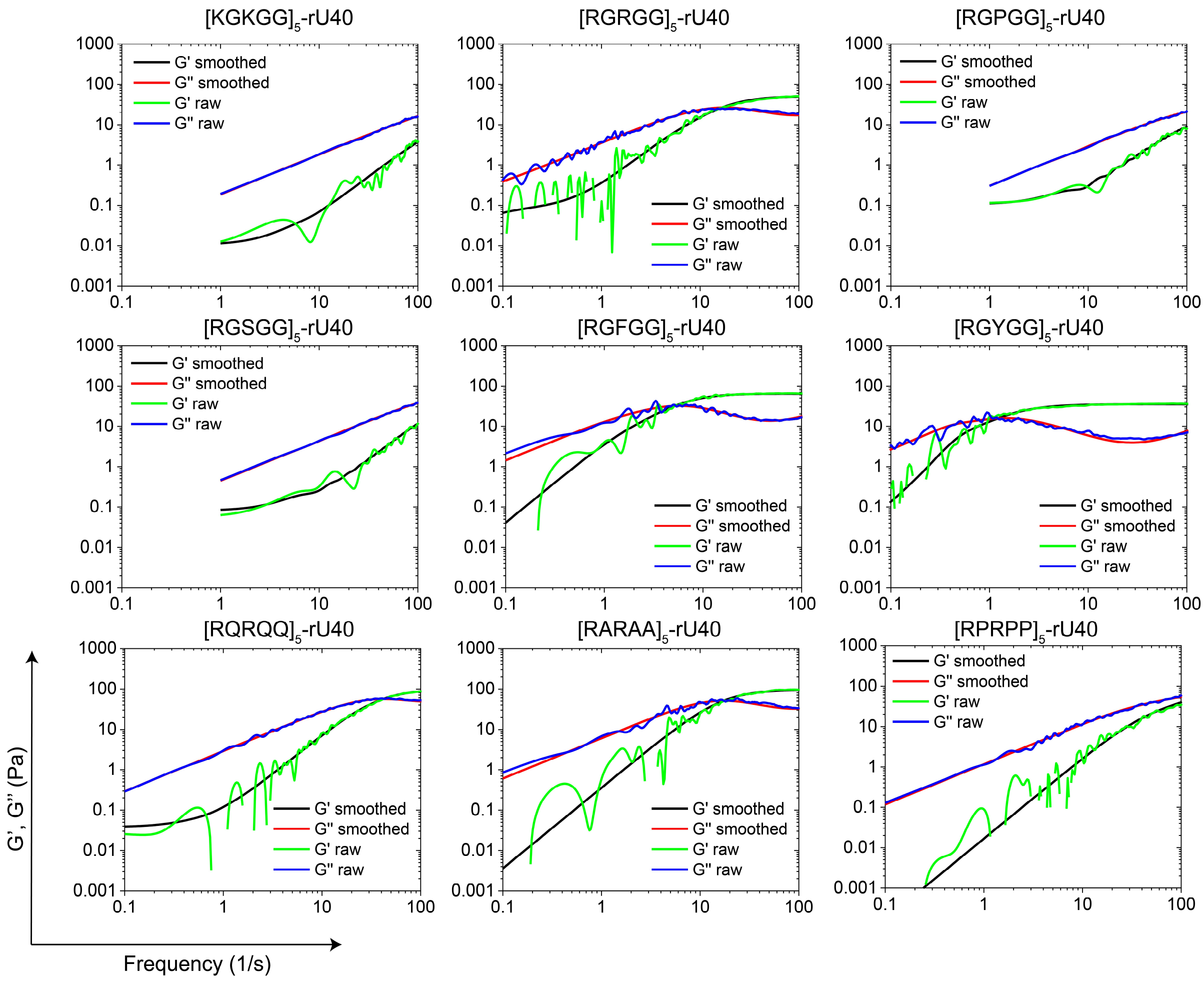


**Fig. S25.** Example plots of the viscoelastic moduli calculated from raw (blue and green) and fitted (black and red) autocorrelation functions (NPAF) of various peptide-NA condensate systems that are shown in Figs. 1-2 in the main text.
